# Human placental stem cell-derived extracellular vesicles ameliorate loss of intestinal stem cell subtypes in experimental necrotizing enterocolitis

**DOI:** 10.64898/2026.11.12.766310

**Authors:** Maryssa A. Ellison, Anchala Guglani, Samuel E. Pomper, Michelle Ebu, Victoria G. Weis

## Abstract

**Background:** Necrotizing enterocolitis (NEC) is a devastating intestinal disease in premature infants with epithelial barrier disruption and loss of intestinal stem cells (ISCs). While depletion of active ISC (aISC) populations has been reported in NEC, the impact of NEC on reserve ISC (rISC) populations during early postnatal development remains poorly understood. We examined developmental and disease-associated changes in ISC subpopulations and evaluated the effects of human placental stem cell-derived extracellular vesicle (hPSC-EV) therapy on ISC preservation and recovery.

**Methods:** Experimental NEC was induced in neonatal rat pups through exposure to NEC stressors with animals assigned as breastfed controls (BF), NEC, or NEC treated with hPSC-EVs (NEC-EV). Ileal tissue was assessed histologically and immunostained for SOX9, OLFM4, HOPX, BMI1, and Ki67. ISC populations were quantified across early postnatal development (32, 56, and 96 hours of life), during NEC progression, and with EV therapy.

**Results:** NEC caused significant histological injury accompanied by marked reductions in SOX9+ and OLFM4+ cells, indicating early and sustained loss of aISC populations. Among rISC markers in established NEC, BMI1+ cells were preserved, whereas HOPX+ cells were significantly reduced. Assessment of healthy early postnatal development revealed that SOX9+ and OLFM4+ cell numbers remained relatively stable, while HOPX+ and BMI1+ populations significantly expanded and were predominantly proliferative. Early NEC did not initially deplete HOPX+ cells but prevented their normal developmental expansion, resulting in the observed reduction of HOPX+ cell abundance during established disease. Treatment with hPSC-EVs significantly improved intestinal architecture and restored SOX9+, OLFM4+, and HOPX+ cell populations to near-control levels.

**Conclusions:** NEC differentially affects ISC subpopulations in the developing intestine, causing loss of aISC and disrupting the expansion of HOPX+ rISC. Preservation and restoration of both aISC and HOPX+ rISC populations may contribute to epithelial regeneration and recovery. These findings identify HOPX-associated development and regeneration as potential therapeutic targets and demonstrate that hPSC-derived extracellular vesicles support ISC recovery and intestinal repair in experimental NEC.

## Introduction

Necrotizing enterocolitis (NEC) remains a significant cause of morbidity and mortality in preterm infants. The incidence of NEC is reported to be as high as 11% among very low birth weight infants (<1500 g) and up to 22% among extremely low birth weight infants (<1000 g)(Hu et al., 2024). Mortality rates remain substantial, ranging from 20% to 30%(Fanaroff et al., 2007; Fitzgibbons et al., 2009; Horbar et al., 2012), with even higher rates observed in infants requiring surgical intervention(Battersby et al., 2018). Recent reports estimate 20-50% of infants with NEC require surgery(S. H. Kim et al., 2025; van Varsseveld et al., 2025), making it the most common neonatal surgical emergency(Akre et al., n.d.). Research over the past decades have identified multiple contributing factors that are involved in NEC pathogenesis, including the loss of the intestinal stem cells (ISCs) that give rise to the epithelial barrier(Barker, 2014; Hosfield et al., 2022; Weis et al., 2021). However, little is understood about the dynamic balance between the different subtypes of ISCs during early life development, disease onset, and active NEC disease. Understanding the mechanisms of epithelial barrier maintenance will yield insights into potential targets and development of therapeutic options to support epithelial regeneration during NEC pathogenesis.

One known contributing factor to epithelial barrier damage in NEC is the increase in cell death, including apoptosis and necroptosis. This cell loss leads to intense break down of the intestinal epithelium when paired with the additional loss of cells within the epithelial crypts, where the ISCs reside(Jilling et al., 2004; Leaphart et al., 2007; Weis et al., 2021; Werts et al., 2019). These ISCs in the intestinal crypts give rise to the differentiated epithelial cell types that are responsible for epithelial function and barrier protection. The current understanding is that there are two subtypes of ISCs that can be generally classified as active stem cell (aISC) or reserve stem cell (rISC). In well-developed adult intestine tissue aISCs are under ongoing proliferation and actively contribute to the replenishment of the epithelial layer under healthy conditions(Barker et al., 2007; Bjerknes & Cheng, 1999; Winton & Ponder, 1990). Locationally, aISCs are found interspersed between Paneth cells within the crypt-base region. Conversely, rISCs sit higher in the +6 / +4 cell position of the crypt. In the healthy adult intestine, the rISCs maintain a low proliferative/quiescent state(Takeda et al., 2011) and are considered resistant to multiple types of damage(Bankaitis et al., 2018a; Gonzalez et al., 2019; Richmond et al., 2017; Roche et al., 2015). rISCs can shift out of a quiescent state and transition into an active proliferative state under intestinal stress and damage to help replenish the lost aISC population(Karmakar et al., 2020; C.-K. Kim et al., 2017; Rees et al., 2020a). Established markers for the aISC population include LGR5 and OLFM4(Flier, Haegebarth, et al., 2009; Yan et al., 2012), while markers of rISCs include, but are not limited to, BMI1, HOPX, mTert, Dclk1, Lrig, and Clu(Bankaitis et al., 2018b). Using a subset of these markers, we can assess changes in the traditionally classified active and reserve ISC subtypes during NEC onset, progression, and repair in the premature intestine.

Previous studies from our group and others have demonstrated the significant decrease of intestinal stem cells in both experimental and clinical NEC ileal tissues(Hosfield et al., 2022; Li et al., 2019; Weis et al., 2021). Our research has illustrated the loss of SOX9 expressing crypt cells in NEC(Weis et al., 2021), a marker that is found broadly expressed on both aISC and rISC populations(Roche et al., 2015; Shi et al., 2013). Furthermore, we and others have also shown the specific loss of aISCs in NEC(Hosfield et al., 2022; Leaphart et al., 2007; Weis et al., 2021). However, there are few studies that have investigated rISCs in NEC pathogenesis. Notably, one study observed that the rISC marker BMI1was preserved in NEC tissue when compared to healthy control tissue(Hosfield et al., 2022), but other rISC markers were not assessed in this study. In various other intestinal injuries, ISC subpopulations have been identified as potential therapeutic targets(Rees et al., 2020b), suggesting protection against ISC loss or support of ISC regeneration may improve NEC outcomes. Thus, understanding the dynamic changes of both active and reserve ISCs during NEC damage and repair is essential for improving protective and therapeutic approaches in the premature intestine.

In addition to examining ISCs in NEC disease, it is important to consider these populations within the context of the developing immature intestine setting, as NEC predominately occurs in preterm infants with developmentally immature gastrointestinal tracts. The expression and developmental dynamics of ISC populations during intestinal development in early postnatal life is not completely understood. Characterizing the ontogenetic shifts in this developmental window of the premature intestine is critical as this period constitutes the baseline environment vulnerable to NEC disease. Similar to the characterization of aISC fluxes, current knowledge of intestinal cell patterns for rISC markers, such as HOPX and BMI1, is derived from studies predominately focused on adult or juvenal tissue and various injury models in mature intestine(Gonzalez et al., 2019; Sangiorgi & Capecchi, 2008; Stewart et al., 2021; Takeda et al., 2011; Wang et al., 2019; Zhang et al., 2026), with few investigations focusing on the fetal or early postnatal timeframe(Creff et al., 2023; Fawkner-Corbett et al., 2021). In this study, we investigated the aISC marker OLFM4 and the rISC markers HOPX and BMI1 to establish their normal development patterns in early-life intestinal development of the premature infant and the subsequent alterations in NEC pathogenesis.

Recent studies have demonstrated the promising therapeutic potential of stem cell-based therapies for NEC, in part by supporting repair of the epithelial barrier and specifically the ISC niche.(Weis et al., 2021) Thus, we also examined the expression of key aISC and rISC markers during tissue repair following treatment with a human placenta stem cell-derived extracellular vesicle therapy. By tracking these active and reserve ISC populations across NEC injury and resolution, we aim to uncover the regenerative cellular programs that guide development, disease pathogenesis, and tissue repair in the premature intestine. Ultimately, this work lays the foundation to identify signaling pathways involved in the epithelial pathophysiology of NEC disease and repair, with the overarching goal to design targeted, developmentally tailored strategies for clinical interventions.

## Methods

### Placental Stem Cell Isolation, Expansion, and Culture

Under an approved protocol by the Institutional Review Board of Wake Forest University School of Medicine, consent was obtained from patients prior to delivery for the collection and use of full-term human placenta specimens. Following placental tissue collection, processing was performed by the Manufacturing Development Center (MDC) within the Wake Forest Institute for Regenerative Medicine (WFIRM), Wake Forest University School of Medicine. As previously described(Weis et al., 2021), chorionic tissue was digested for the isolation of perinatal placental cells and stocks of primary human placental stem cells (hPSC) were cryopreserved for subsequent use following expansion and c-kit positive selection. Cells were thawed and plated in tissue culture flasks with α-MEM media (HyClone, Logan, UT), 10% fetal bovine serum (FBS), and 1× Penicillin-Streptomycin. Cells were cultured at 37°C, 5% CO2, and no more than 80% confluency. Media was exchanged for media containing extracellular vesicle-depleted FBS and incubated for 48 hours. The 48-hour conditioned media (CM) was collected and centrifugated at 300 × g for 10 minutes to remove cell debris. CM supernatant was collected to not disturb any cell debris pellet, transferred to sterile polystyrene bottles, and placed in −20°C until further processing (less than 6-months storage).

### Placental Stem Cell Extracellular Vesicle Isolation and Characterization

CM was further processed for hPSC extracellular vesicle (EV) isolation by thawing CM, ultrafiltration using Amicon Stirred Cell 100kD ultrafiltration discs (MilliporeSigma, Burlington, MA), and concentration using Amicon Ultra-15 Centrifugal Filter Units (MilliporeSigma, Burlington, MA).

Following filtration and concentration, 500 *μ*l of condensed CM is processed via size exclusion chromatography using qEV2 / 35 nm Gen 2 columns (Izon Science, Medford, MA) to achieve high concentrations of small EV isolations within a recovery range of 35-350 nm. Nanoparticle tracking analysis (NTA) (NanoSight Pro, Malvern Panalytical, Westborough, MA) was utilized to determine the size and concentrations of hPSC-EV isolations using the following parameter settings: blue 405 laser in scatter mode, camera level 12 (NTA 3.0 levels), shutter of 600, slider gain of 350, 25 frames per second at a total of 1499 frames captured, detection threshold of 5, and max jump distance at auto 12.9-13.9 pixels. Presence of transmembrane tetraspanin proteins (CD9, CD81, CD63) was confirmed using the ONI EV Profiler 2 Application Kit with stochastic optical reconstruction microscopy on a Nanoimager (ONI, San Diego, CA). EV samples were prepared for imaging as outlined in the kit protocol. Transmission electron microscopy (TEM) was used to confirm EVs within the desired size range and expected EV morphology. These EV characterizations of size, morphology, concentration, and tetraspanin expression are in accordance with MISEV2018 guidelines described by the International Society for Extracellular Vesicle(Théry et al., 2018).

### NEC Animal Model

All animal procedures were performed under protocols approved by the IACUC of Wake Forest University School of Medicine. Three experimental groups of newborn Sprague-Dawley rats were utilized for this study: healthy breastfed control (BF), NEC disease (NEC), and NEC with hPSC-EV therapy (NEC-EV). The BF group remained with dam throughout the entirety of the study. The NEC and NEC-EV groups were isolated from dams within 6 hours of birth, housed in a temperature (32°C) and humidity (50%) controlled incubator, and underwent NEC induction through exposure to three factors known to trigger and perpetuate the NEC-like disease state, as previously described(Weis et al., 2021). Briefly, pups were orally gavaged every 6 hours with hyperosmolar formula prepared from a ratio of Similac/Esbilac (15 g Similac Advance (Abbott Nutrition, Columbus, OH) in 75 mL Esbilac canine supplement (Pet-Ag, Hampshire, IL))(Weis et al., 2021). In addition, pups received three doses of oral lipopolysaccharide (LPS, Cat no. L3012, Sigma-Aldrich, St. Louis, MO) within 30 hours of birth with formula feeds (4 µg/g for each dose). Pups also underwent whole-body hypoxia induction for 10 minutes, three times per day, with hypoxic conditions of 4-6% oxygen. 32 hours after birth, NEC-EV pups received a first dose of hPSC-EV therapy at a concentration of 1×10^10^ EVs in a 50 *μ*l intraperitoneal injection (IP), with a second dose administered at 56-hours after birth (24-hours after first dose) for a total of 2×10^10^ EVs per pup. The NEC group received 50 *μ*l sham injection of saline (diluent for hPSV-EV) at the same time points. Pups were maintained and monitored for humane endpoints per IACUC protocol standards. At 96 hours of life (day 4), all pups were humanely euthanized and immediately processed for tissue harvesting.

### Tissue Harvest and Processing

The entire intestine, duodenum to the distal colon, was collected and washed with 1× phosphate-buffered saline (PBS). Tissue underwent overnight fixation in 4% paraformaldehyde at 4°C followed by transfer to 70% ethanol prior to routing processing and embedding in paraffin blocks. Tissue blocks were sectioned at 4 µm thickness, mounted to coated glass slides, air-dried overnight, and stored at room temperature until further use.

### Histological Assessment and NEC Grade

Sections were deparaffinized and rehydrated for hematoxylin and eosin (H&E) staining. An established scoring system was used to assess NEC damage and a NEC Grade ranging from 0 (normal) to 4 (transmural necrosis) was assigned based on analysis of H&E-stained sections that included the full ileal length, as previously described (Weis et al., 2021). NEC damage is defined as grade 2 or above. Representative images of H&E-stained sections were imaged at 20× using an EVOS FL Auto 2 Cell Imaging System (Invitrogen, CA).

### Immunostaining and Quantifications

Paraffin sections were heated at 60°C for 30 minutes and cooled to room temperature followed by routine deparaffinization and rehydration. For antigen retrieval, sections were submerged in 1× Target Retrieval Solution, Citrate pH 6.1 (Agilent DAKO Cat No. S169984-2) and placed in a pressure cooker for 15 minutes on high heat setting. After cooling on ice and washing with deionized water and 1× PBS, sections were blocked with Protein Block, Serum-Free (Agilent DAKO Cat No. X090930-2) for 1 hour at room temperature in a humidity chamber. The following primary antibodies were incubated overnight at 4°C in a humidity chamber: Rabbit anti-OLFM4 (1:1000; Cat No. 39141T, Cell Signaling Technologies, Danvers, MA), Rabbit Anti-SOX9 (1:1000; Cat No. AB5535, Sigma, St. Louis, MO), Mouse anti-Ki67 (1:50; Cat. No 550609, Waters Biosciences, Milford, MA), Rabbit Anti-Ki67 (1:750; Cat. No AB16667-1001, Abcam, Waltham, MA), Rabbit Anit-BMI1 (1:75; Cat No. 6964T, Cell Signaling Technologies, Danvers, MA), Mouse Anti-HOPX/Hop E-1 (1:100; Cat No. sc-398703, Santa Cruz Biotechnology, Dallas, TX). After three 5-minute washes of 1× PBS, sections were incubated at room temperature for 1 hour with secondary antibodies conjugated for immunofluorescence: Anti-Rabbit Alexa Fluor 488 (1:1000; Jackson ImmunoResearch, PA), Anti-Mouse Alexa Fluor 488 (1:1000; Jackson ImmunoResearch, West Grove, PA), Anti-Rabbit Alexa Fluor 594 (1:1000; Jackson ImmunoResearch, PA), Anti-Rabbit CF-594 (1:1000; Biotium, Fremont, CA). Sections were then washed three times with 1× PBS. Nuclei were stained with 10mM 40,6-diamidino-2-phenylindole (DAPI; 1:10,000) for 5 minutes. After a final washing sequence, sections were mounted with ProLong Gold Antifade Reagent (Invitrogen, Carlsbad, CA).

Sections were imaged using a Leica DM4000B fluorescent microscope. For each animal, three images were taken along the length of the ileum. Well-oriented crypt/villus units and identifiable lumen space were noted prior to image capture. For each animal, OLFM4, SOX9, HOPX, BMI1, and Ki67 immunostainings were quantified from these three randomly selected images, which was approximately 700 µm of total tissue length at 20× magnification. Manual cell count quantification was performed to identify positively stained cells within the crypt region utilizing ImageJ 1.53t. Cell counts were normalized by tissue length quantified (per millimeter of ileum length assessed).

### Statistics

Results are expressed as mean ± standard deviation for each analysis with statistical testing using GraphPad Prism v10.4.2. For comparisons between 2 groups (e.g., BF-32h vs NEC-32h), two-tailed Student’s T-test was performed. For comparison of three groups (e.g., BF-32h vs BF-56h vs BF-96h), one-way ANOVA testing with Bonferroni’s post-hoc correction was performed. The *α*-level for statistical significance was set at p<0.05 for all statistical tests.

## Results

### hPSC-EV Therapy Ameliorates NEC Histological Damage

To first confirm therapeutic impact to overall histological tissue damage, we examined H&E-stained ileal sections from BF, NEC, and NEC-EV groups (Figure 1). BF control pups showed normal ileal morphology with no histological damage (NEC grade 0). In contrast, NEC pups showed significant epithelial disruption with loss of crypt-villus morphology and cellular sloughing, consistent with NEC histological damage scores (grade 2 or above). hPSC-EV treatment significantly ameliorated NEC damage, with reemergence of normal crypt-villus architecture and reduced epithelial damage, consistent with NEC grade 1 and representing a significant decrease in NEC grade.

**Figure 1.**
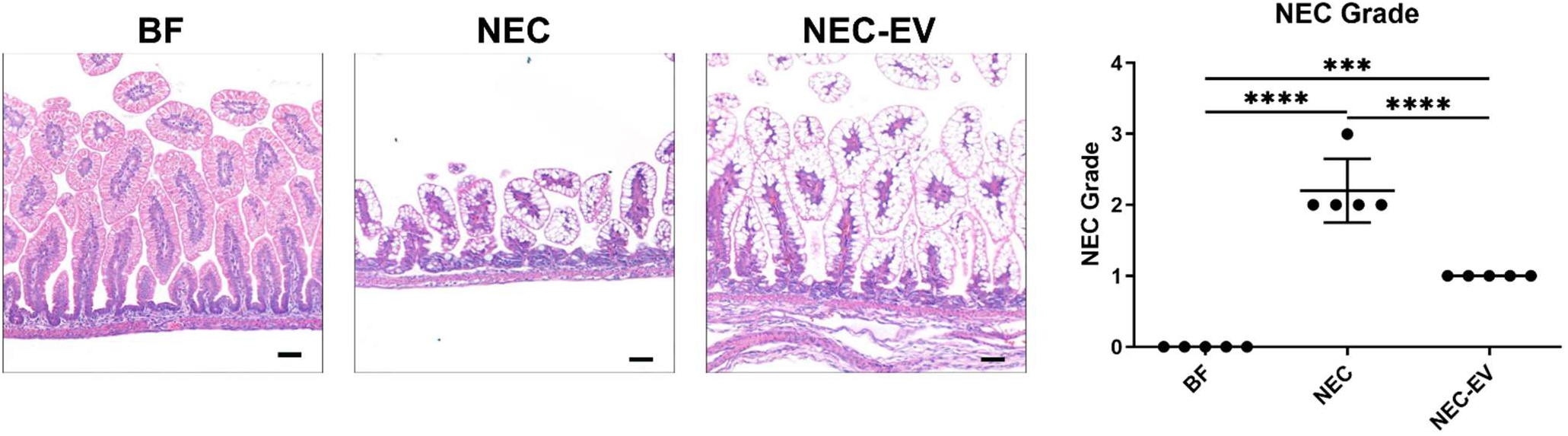
H&E-stained ileal sections from BF, NEC, and NEC-EV show significant loss of crypt/villus morphology and villi blunting in NEC. NEC treated with hPSC-EV has partial recovery of the crypt/villus structures. Scale bar = 50 μm, \*\*\**p<*0.001, \*\*\*\**p<*0.0001, BF, breastfed; NEC, necrotizing enterocolitis; NEV-EV, necrotizing enterocolitis with hPSC extracellular vesicle therapy.

### NEC Pathogenesis Reduces Both Active and Reserve Intestinal Stem Cells and is Ameliorated with hPSC-EV Therapy

We next assessed the broad ISC marker of SOX9 (SRY-box transcription factor 9), which is known to be expressed by both active and reserve intestinal stem cell populations at varying levels(Bankaitis et al., 2018a). In established experimental NEC disease at 96 hours, SOX9+ cells were significantly reduced compared to healthy BF controls (BF: 338.57±54.68 cells/mm, NEC: 146.23±39.85 cells/mm, *p*<0.001) (Figure 2A). This SOX9+ cell loss in NEC was improved with hPSC-EV therapy (NEC-EV: 285.09±52.47 cells/mm, *p*<0.01). Next, we quantified OLFM4+ cells to specifically investigate aISCs in NEC and repair. Similar to SOX9, OLFM4+ cells were lost in NEC and were recovered with hPSC-EV treatment (BF: 72.98±16.85 cells/mm, NEC: 35.72±16.07 cells/mm, NEC-EV: 66.61±7.74 cells/mm BF vs NEC: *p*<0.01, NEC vs NEC-EV: *p*<0.05). (Figure 2B).

**Figure 2.**
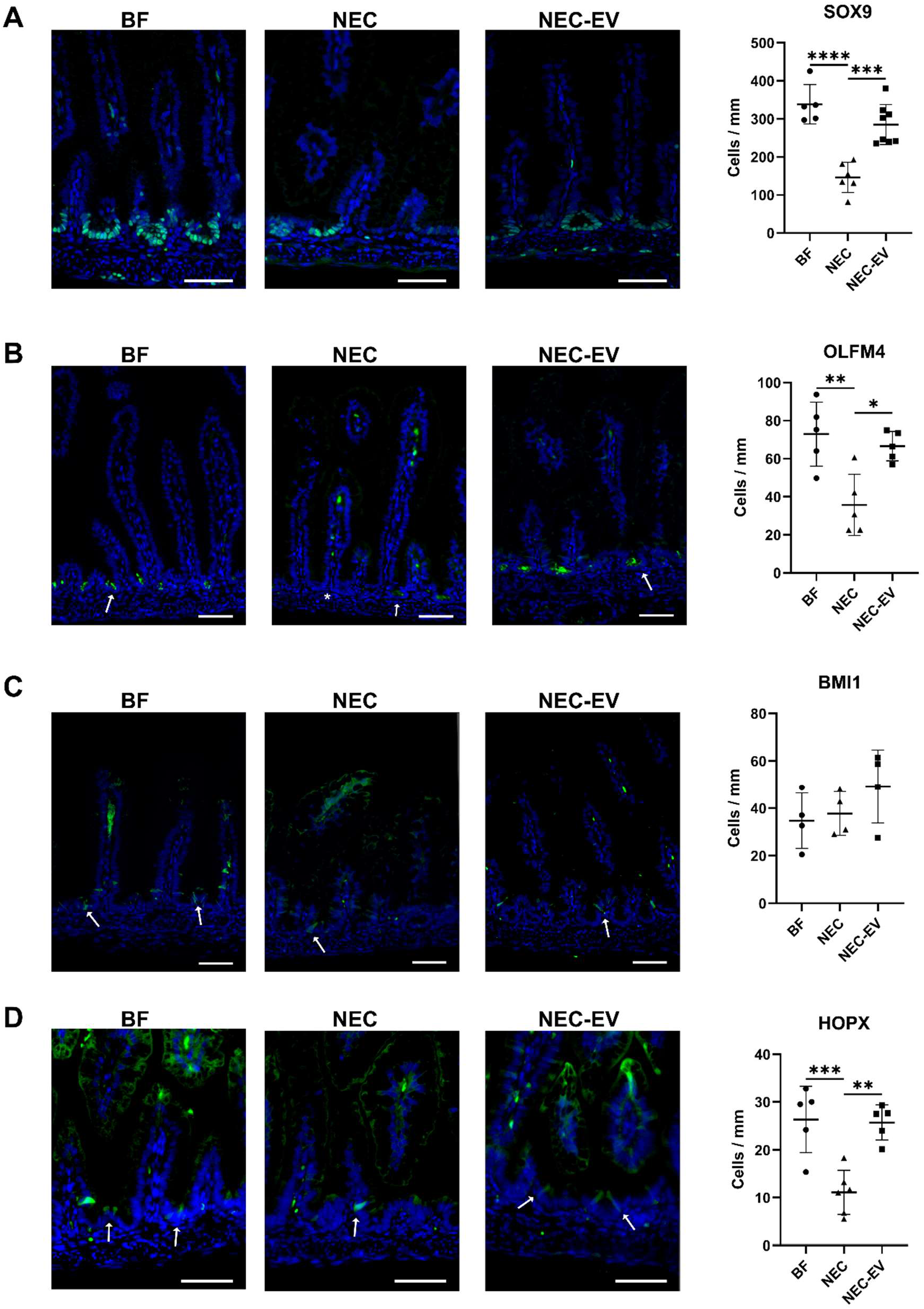
Assessment of active and reserve stem cell markers within the crypts in BF, NEC, and NEC-EV groups at 96 hours of life (day 4). (A) Immunostaining (SOX9 = green, DAPI = blue) demonstrates a high presence of SOX9 in BF crypt region, with a loss seen in NEC crypts and reemergence with NEV-EV therapy. Quantification of SOX9+ cells shows a significant decrease in NEC and a significant increase with hPSC-EV therapy. (B) Immunostaining (OLFM4 = green, DAPI = blue) demonstrates crypt presence of OLFM4+ cells (arrows), with a decrease noted in NEC (asterisk). Quantification of OLFM4+ cells shows a significant decrease in NEC with a significant increase in NEC-EV. (C) Immunostaining (BMI1 = green, DAPI = blue) demonstrates BMI1-positive cells both within the crypt region (arrows) and occasionally in the transit-amplifying region. Quantification of BMI1+ cells in crypt regions shows no changes across the cohorts. (D) Immunostaining (HOPX = green, DAPI = blue) demonstrates HOPX+ cells within the crypt region (arrows), while also noting infrequent HOPX+ cells along the villi in the transit-amplifying region. Quantification HOPX+ cells in crypt regions shows a significant decrease in NEC, with a significant increase in NEC-EV. Quantifications are expressed as mean ± standard deviation of cell counts normalized per length of tissue analyzed. Samples sizes for each analysis are denoted in dot scatter plots, with n<4 pups per group. Scale bar = 50 µm, \**p<*0.05, \*\**p<*0.01 \*\*\**p<*0.001, \*\*\*\**p<*0.0001.

In contrast to aISC, rISC populations are traditionally thought to be resistant to damage. Thus, we next investigated the classically categorized rISC populations via the two rISC markers of BMI1 and HOPX. Compared to healthy BF controls, BMI1 remained unchanged in active NEC as well as in NEC with EV therapy (Figure 2C). Notably, a previous study also observed no changes in BMI1+ cells during NEC pathogenesis(Hosfield et al., 2022). While BMI1+ cells have a slow proliferation rate in mature intestine, recent studies have shown that BMI1+ cells can expand during early life development of the intestine (Dehmer et al., 2011; Smith et al., 2025) and play an important role in establishing the mature ISC populations. Accordingly, we found that almost all BMI1+ cells at Day 4 of life were also Ki67+ (data not shown) and no significant differences in proliferating BMI+ cells were observed between groups. These data suggest BMI1+ cells, although proliferative, are resistant to NEC damage. We next assessed HOPX+ cells as a rISC marker known to play roles in early intestinal development and regeneration following damage. In NEC disease, HOPX+ cells were significantly reduced (BF: 26.39±6.89 cells/mm, NEC: 11.11±4.63 cells/mm, *p*<0.001) (Figure 2D). Similar to aISC loss, HOPX+ ISC loss in NEC was ameliorated with EV therapy (NEC-EV: 25.75±3.64 cells/mm, *p*<0.01) (Figure 2D). Quantification of the proliferating and non-proliferating subsets of HOPX+ cells revealed a significant decrease in proliferating HOPX+ cells (BF: 21.99±7.92 cells/mm, NEC: 6.90±2.53 cells/mm, *p*<0.001) while non-proliferating HOPX+ cells were unchanged in NEC (Figure 3). Furthermore, NEC-EV pups exhibited a significant increase in proliferating HOPX+ cells compared to NEC group (NEC-EV: 18.97±3.25 cells/mm, *p*<0.001) (Figure 3). Together, these data establish loss and recovery of aISC and a potential unique role of HOPX+ rISC in NEC pathogenesis and repair.

**Figure 3.**
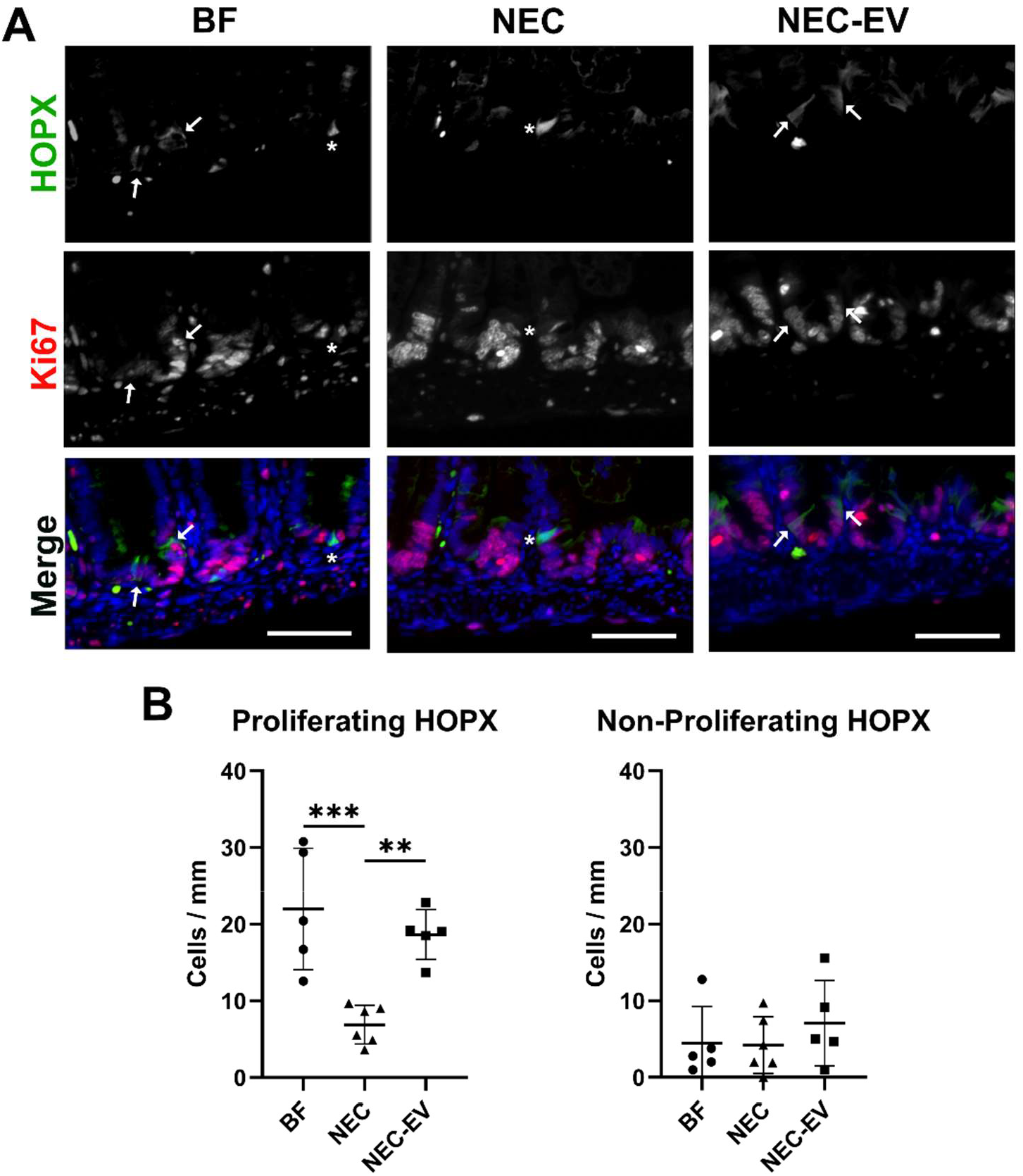
Proliferating and non-proliferating HOPX+ cells in BF, NEC, and hPSC-EV therapy. (A) Immunostaining of HOPX (green), Ki67 (red), and DAPI (blue) demonstrates HOPX/Ki67-dual positive cells (arrows, Hopx+Ki67+) and HOPX only-positive (asterisks, HOPX+Ki67-) at 96 hours. (B) Quantification of proliferating HOPX+ rISCs (HOPX+Ki67+) in the crypt regions shows a significant decrease in NEC with a significant increase upon NEC-EV therapy. Quantification of non-proliferating HOPX+ rISCs (HOPX+Ki67-) in the crypt revealed no significant changes. Quantifications are expressed as mean ± standard deviation of cell counts normalized per length of tissue analyzed. Samples sizes for each analysis are denoted in dot scatter plots, with n=5-6 animals per group. Scale bar = 50 *μ*m, \*\**p<*0.01 \*\*\**p<*0.001.

### HOPX+ Cells, but not SOX9+ or OLFM4+ Cells, Increase during Early Life Development

As we saw the majority of HOPX+ cells were proliferative with Ki67-positivity at Day 4 in healthy BF controls, we sought to establish the normal early life trajectory of the ISC populations found aberrant in NEC disease. SOX9+ cells were consistently stable from 32 hours of life to the 96-hour (Day 4) endpoint (Figure 4A). OLFM4+ cells also did not have any significant changes in early life in healthy BF ileum, though a slight trend upward may suggest further expansion past 96 hours (Figure 4B).

**Figure 4.**
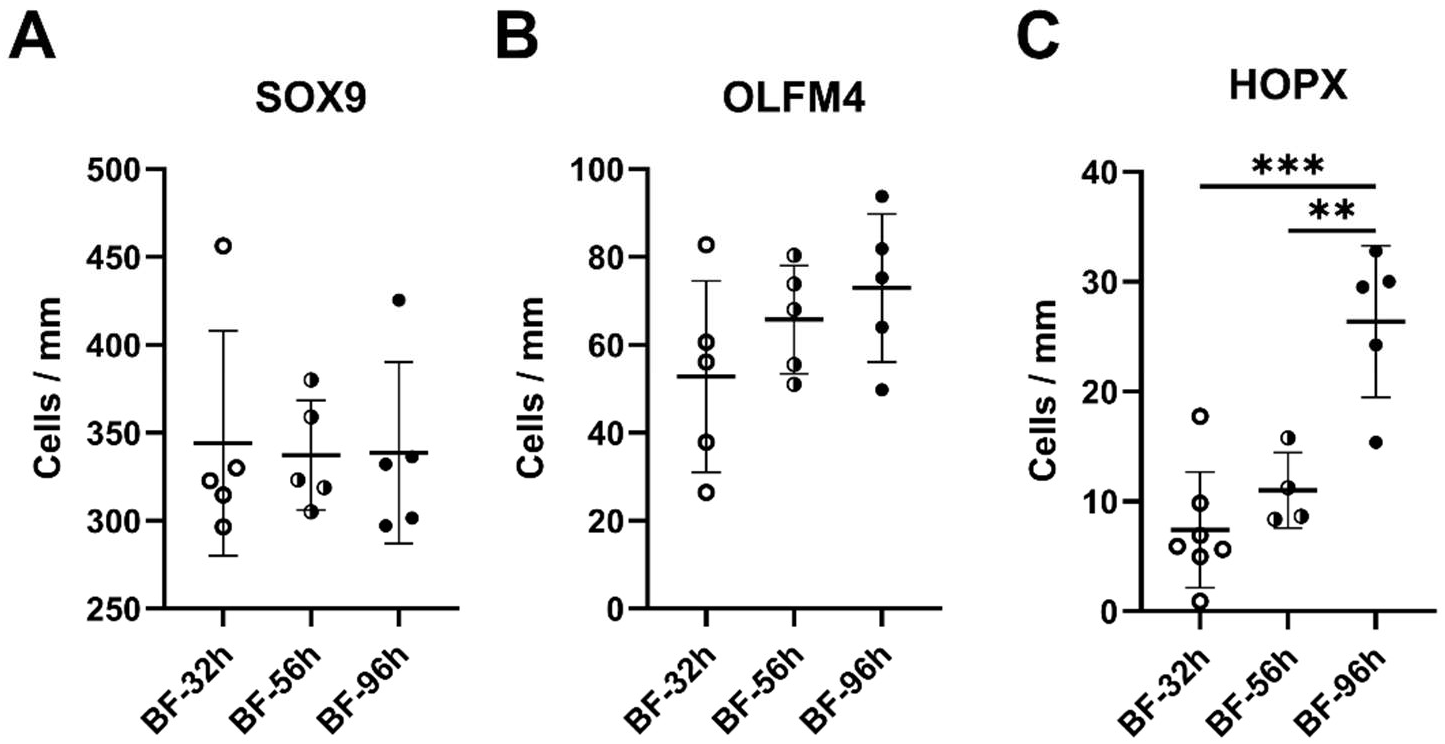
Developmental time course assessment of the intestinal stem cell markers SOX9, OLFM4, and HOPX in BF ileum. (A) SOX9+ cell abundance was observed in steady state from 32 hours to 96 hours. (B) OLFM4+ cell abundance was observed with a slightly increasing trend from 32 hours to 96 hours. (C) HOPX+ cell abundance was observed to significantly increase in early healthy postnatal development at 96 hours as compared to 32 hours and 56 hours. Quantifications are expressed as mean ± standard deviation of cell counts normalized per length of tissue analyzed. Samples sizes for each analysis are denoted in dot scatter plots, with n=4-7 animals per group. \*\**p<*0.01 \*\*\**p<*0.001.

Conversely, a significant increase in HOPX+ cells occurred at 96 hours of healthy development compared to both the earlier 32-hour and 56-hour timepoints (BF32h: 7.43 ±5.27 cells/mm vs 96h *p*<0.0001, BF56h: 11.01±3.43 cells/mm vs 96h *p*<0.01) (Figure 4C). As shown in Supplemental Figure 1A, the majority of these HOPX+ cells within each timepoint are also Ki67+, suggesting they are in active cell proliferation cycle. Following this increasing trend in traditionally classified HOPX+ rISC, we compared the number of BMI1+ cells at the earliest time point of 32 hours of life to 96 hours.

Similar to HOPX+ cells, there was a significant increase in the total number of BMI1+ cells at 96 hours of life and they were predominately also Ki67+ (Supplemental Figure 1B). Together, these data demonstrate an active expansion of HOPX+ and BMI+ ISC populations in early healthy development, while SOX9+ and OLFM4 + cell numbers remain consistent.

### NEC Pathogenesis Causes Early Loss of SOX9+ and OLFM4+ Cells Coupled with a Blockade of the Normal Expansion of HOPX+ Cells

To better understand the impact of NEC disease on the ISC niche, we investigated whether early NEC damage reduced the pool of each ISC population or rather blocked the natural healthy developmental expansion of specific populations such as the HOPX+ and BMI1+ cells. As shown in Figure 5A-B, there is a significant loss of both SOX9+ and OLFM4+ cells in the earliest stages of NEC damage at 32 hours (SOX9: BF32h: 344.1±6.97, NEC32h: 181.2 ± 18.94, *p*<0.001) (OLFM4: BF32h: 52.80 ± 21.72,

**Figure 5.**
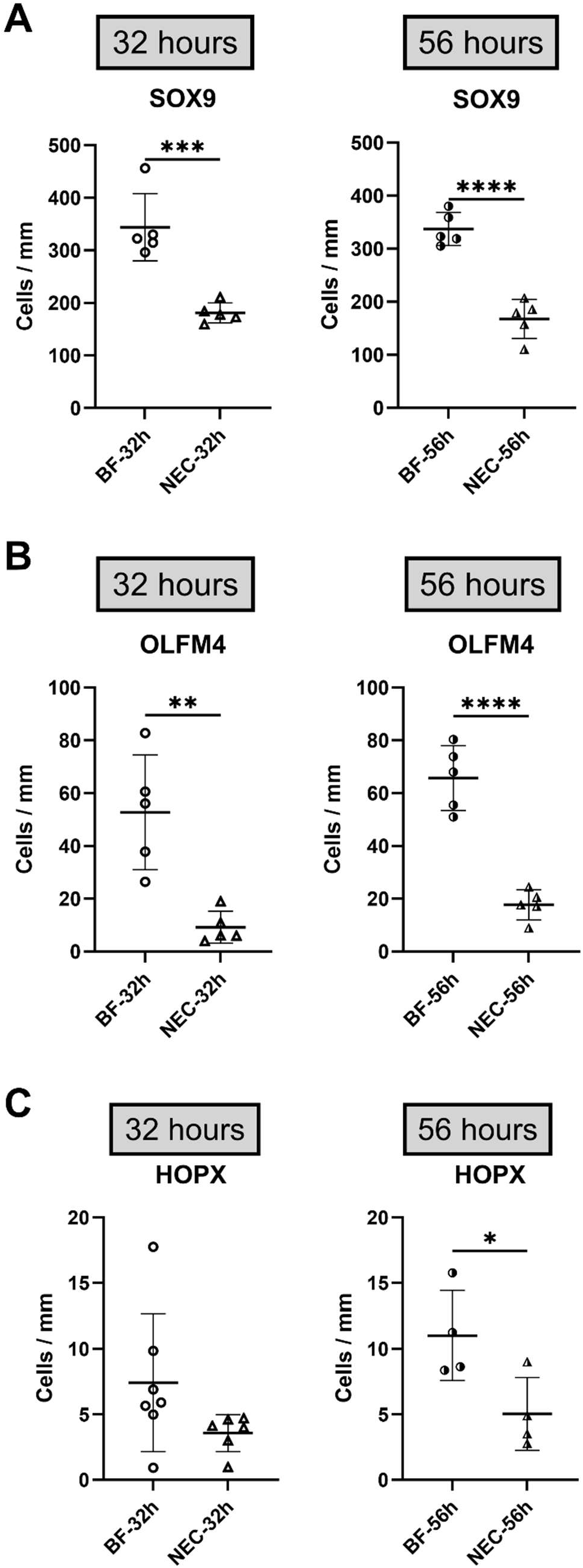
Comparison of the intestinal stem cell markers SOX9, OLFM4, and HOPX in early-stage NEC damaged ileum. (A) SOX9+ cells are significantly decreased in early NEC damage at both 32-hours and 56-hours as compared to BF controls. (B) OLFM4+ cells are significantly decreased in early NEC damage at both 32-hours and 56-hours as compared to BF controls. (C) No significant changes are found in HOPX+ cells in early NEC at 32 hours, while HOPX+ cells are significantly decreased in NEC at 56 hours as compared to BF controls. Quantifications are expressed as mean ± standard deviation of cell counts normalized per length of tissue analyzed. Samples sizes for each analysis are denoted in dot scatter plots, with n=4-7 animals per group. \**p<*0.05, \*\**p<*0.01 \*\*\**p<*0.001, \*\*\*\**p<*0.0001.

NEC32h: 9.23± 6.05, *p*<0.01). However, there is no change in HOPX+ cells in early NEC at 32 hours (Figure 5C). These data suggest initial damage to the vulnerable aISC populations during early NEC onset. A similar reduction of SOX9+ and OLFM4+ cells was also observed at 56 hours of NEC damage compared to healthy BF controls at 56 hours of NEC damage (SOX9: BF56h: 337.30±31.11, NEC56h: 167.77±36.65, *p*<0.0001) (OLFM4: BF56h: 65.77±12.30, NEC56h:17.77 ±5.72, *p*<0.0001) (Figure 5A and Figure 5B). Notably though, the number of SOX9+ and OLFM4+ cells in NEC at 56 hours is similar to that found at the early 32-hour timepoint, suggesting there is not a further reduction of aISC as NEC progresses into established, but not fully necrotic-stage of disease. In contrast, HOPX+ cells that had no change in early NEC onset were found to be reduced as NEC progressed at 56 hours compared to healthy controls (BF56h:11.01±3.43, NEC56h: 5.04±2.78, *p*<0.05) (Figure 5C). However, the number of HOPX+ cells is similar at both 32 hours and 56 hours of NEC, suggesting the significant change at 56 hours is due to the trend of HOPX+ cell expansion in normal healthy development. In distinguishing between proliferating and non-proliferating HOPX+ cells, we found no substantial differences between healthy and NEC ileum at either 32 or 56 hours (Supplemental Figure 2A). Interestingly, while there was no change in the other rISC marker of BMI1+ cells in established NEC disease (Figure 2C), there was a small but significant loss of BMI1+ cells in early NEC at 32 hours (Supplemental Figure 2B).

However, this early loss appeared to be recovered as NEC damage progressed at 96 hours, further suggesting a resistance in this ISC population to NEC damage. This data suggests that while NEC causes an active loss of SOX9+ and OLFM4+ cells, NEC pathogenesis inhibits the normal expansion of HOPX+ cells, rather than a true loss of the present HOPX+ cells (Figure 6). Together, this aISC loss coupled with the inhibition of rISC expansion would leave the immature epithelium vulnerable to further damage and disease progression.

**Figure 6.**
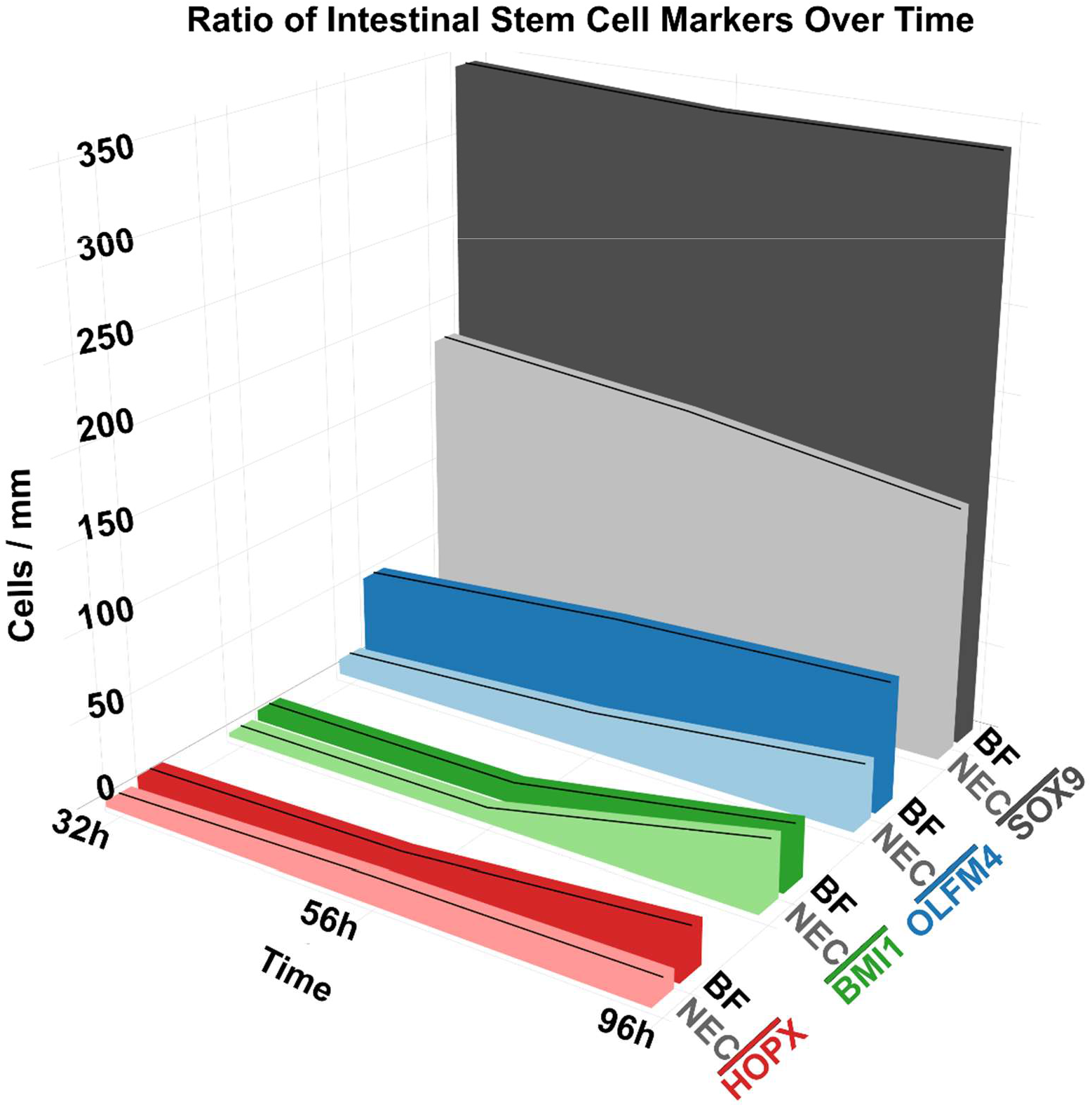
Visual comparison of ISC markers throughout early postnatal development demonstrates relative abundance of SOX9, OLFM4, HOPX, and BMI1 positive cells in ileal tissue at 32-hour, 56-hour, and 96-hour timepoints in both healthy BF and NEC disease. Quantifications are summarized and expressed as mean of cell counts normalized per length of tissue analyzed.

## Discussion

It is critical to consider developmental timing in conjunction with disease pathogenesis when investigating NEC, as intestinal physiology changes significantly throughout early-life developmental stages. In addition to observed physical structural changes, a multitude of signals and factors contribute to the maturation of the intestine and differ throughout developmental stages(Venkatraman et al., 2021). More specifically, key signals acting on ISC development differ in actively maturing tissue as compared to developed tissue in homeostasis. For example, in the developed mature intestine, conical Wnt singling is isolated to the crypt region, while in early fetal and postnatal development this paracrine signal is found along the intestinal villus(Venkatraman et al., 2021). Transcription factor expression also differs based on intestinal developmental stages. SOX9, a downstream target of Wnt signaling, is expressed along the entire epithelium in early embryonic development in mice, while it is expressed only in crypt regions in the adult intestine(Blache et al., 2004). Therefore, it is important to re-examine what is considered “dogma” for ISCs and ISC subtypes in early developmental studies, as much of the current evidence is based on expression patterns determined in fully developed adult tissue. Our NEC model accounts for developmental variation by employing full gestation neonate rat as our model system because neonate rat intestine is comparable to that of a human fetus in the late second or early third trimester(Drozdowski et al., 2010). This developmental staging is consistent with clinical NEC disease presentation that occurs predominantly in the preterm infant population(Shulhan et al., 2017). NEC pathogenesis research therefore should consider the dynamic developing preterm intestine as a critical determinant of disease pathology when interpreting preclinical findings and translational applicability.

There is limited data on SOX9 expression in neonates(Venkatraman et al., 2021), but it is identified as crucial for intestinal development as a transcriptional regulator for epithelial cell differentiation and essential for Paneth and goblet cell formation(Bastide et al., 2007). SOX9 is also linked to proliferative capacity, with lower levels associated with an increased proliferative state, and higher levels linked to decreased proliferation. SOX9 thereby aids in both reserve stem cell quiescent capacity and promoting terminal differentiation(Formeister et al., 2009). In this study, we found an abundance of SOX9+ cells present in the crypt region of the villi in healthy control BF controls at the earliest time point (32 hours of life) with maintained steady state presence through 96 hours of life. SOX9+ cells significantly reduced in NEC pups compared to healthy BF pups but are still fairly abundant during disease. We previously found SOX9 to be significantly reduced at 96 hours of life in NEC injury(Weis et al., 2021), and show in this study that SOX9 is significantly reduced as early as 32 hours of NEC injury, with continued significant reduction compared to healthy tissue. Loss of this ISC population in the early stages of NEC progression likely significantly contributes to the progressing severity of damage in the epithelial and extent of overall intestinal damage.

To understand the dynamic relationships of ISC subsets in development and disease, we first investigated OLFM4, a marker of aISCs. OLFM4+ cells maintained a constant presence from 32 to 96 hours of life in healthy tissue with no difference observed during the early developmental timeframe. However, OLFM4 was reduced in NEC at each time point throughout early and established disease. OLFM4 has been considered as a strong marker for LGR5 aISCs(Flier, Gijn, et al., 2009; Flier, Haegebarth, et al., 2009), but recent work suggests that the maker may not be reliable in the strict identification of LGR5+ aISCs in humans(Liu & Rodgers, 2022). Nonetheless, our neonate rat ileal immunostaining for OLFM4 identified positive cells localized to the crypt base region and maintained a heterogenous crypt cell pattern. Given our data, combined with the current lack of anti-rat LGR5 antibodies at this time, OLFM4 is likely to be the most reliable method for aISC identification in rat small intestine. Therefore, the loss of this aISC population within our model potentially accounts for the reduction in native regenerative capacity of the epithelium during injury.

Conversely, the rISC markers, HOPX and BMI1, while low in overall abundance, were observed at increasing quantities during healthy development. Correspondingly, HOPX+ cells were not reduced in early NEC, but compared to the increasing HOPX+ cell numbers in healthy pup development, later stage NEC (56- and 96-hours) did have decreased HOPX+ cells. Previous studies examining HOPX+ cells under a different injury and repair condition have identified their reserve capacity during acute injury, with preserved cell populations in the setting of early pre-acute injury(Stewart et al., 2021). Similarly, in our study, HOPX+ cell populations were preserved in early NEC at 32 hours, however, HOPX+ cells were significantly decreased in established NEC as compared to BF controls. While a slight increasing trend is observed throughout NEC induction, HOPX+ cells are reduced in established NEC relative to their observed expansion in BF controls. These cellular dynamics suggest disruption of the normal healthy developmental increase in HOPX+ cells in early postnatal development under sustained NEC injury stress. Our study also explored an additional reserve stem cell marker, BMI1, with observed increased expression over the course of early postnatal development as previously reported(Hosfield et al., 2022). While our data show a small early loss of BMI1+cells during NEC onset (32 hours), there is no significant reduction of BMI1+ cells in established NEC (96 hours) as compared to developmentally-matched control BF tissue. These non-changes in established NEC are consistent with previously published reports supporting the general resistant nature of BMI1+ cells during injury(Gonzalez et al., 2019; Hosfield et al., 2022; Sangiorgi & Capecchi, 2008; Yan et al., 2012). Our study supports that BMI1 may not be as involved in NEC pathogenesis compared to other ISC subpopulations(Hosfield et al., 2022). However, lower abundance of HOPX+ and BMI1+ rISC compared to aISC compartments may warrant further investigation for their role in NEC development, progression, and repair.

There is a limited understanding of the role of rISC in NEC. These studies of rISCs are particularly as there are no clear single expression profile that can conclusively identify a truly quiescent rISC. Known rISC markers can be variably co-expressed with other rISC markers and at variable expression levels(Bankaitis et al., 2018b). Moreover, these co-expressions can dynamically change depending on the developmental age, injury status, or healing stage. One limitation of this study is that co-expression of our two selected rISC markers was not directly included in our experimental design. We prioritized co-localization with Ki67 to assess proliferation status of each rISC subpopulation. Future studies should more comprehensively investigate co-expression of these markers to identify the ISC phenotypes present during healthy intestinal development and in NEC disease.

Our data suggests a potentially important role for HOPX-expressing rISCs during NEC injury, consistent with published work on HOPX-expressing rISCs in intestinal injury repair with HOPX silencing experiments(Stewart et al., 2021)(Zhang et al., 2026). HOPX expression has been shown as important in maintaining a functional rISC population that can be activated after intestinal epithelial damage to subsequently replenish the LGR5+ aISC compartment(Zhang et al., 2026). Taken together with the results from our study, we propose that amelioration of HOPX+ cell aberrations in NEC may be important for recovery of the intestinal epithelium and long-term intestinal development and health, where it can act as a crucial molecular switch controlling intestinal stem cell proliferative activity(Gonzalez et al., 2019). Long-term studies would be insightful to address this gap but require more technically challenging model due to the limited feasibly timeframe of this NEC model. Future studies could assess the HOPX+ cell population after natural recovery from NEC to identify whether the inhibited expansion alters long-term crypt balance or if this cell population recovers with time.

hPSC-EV therapy ameliorated tissue histological damage. This amelioration was observed in both the aISC and rISC compartments with recovery of SOX9+, OLFM4+, and HOPX+ ISCs. However, it is unknown the exact impact of each ISC compartment recovery has on the overall repair of the histological damage. To identify the impact of the HOPX+ cells versus the OLFM4+ cells and any other ISC subpopulations in NEC recovery, more comprehensive transgenic experimental models are needed. These studies could include cell lineage tracing studies or alternatively cell knockout studies to better assess each compartments impact on NEC recovery. These studies could assess both natural NEC recovery as well as hPSC-EV treatment-induced repair to determine the dependence of therapeutic benefits of hPSC-EV on ISC-mediated regenerative mechanisms.

To continue to progress our understanding of NEC pathogenesis and potential therapeutic repair, studies must investigate NEC disease within the dynamic context of neonatal ISC developmental maturation. By assessing the developmental trajectories in both healthy developing tissue and associated injured developing tissue, researchers can more fully elucidate cellular mechanisms in NEC pathogenesis and delineate true cellular loss versus alterations in normal developmental paths. In this study, we quantified the ISC markers of SOX9, OLFM4, HOPX, and BMI1 throughout early postnatal developmental timepoints in healthy breastfed and NEC-injury induced neonatal rat intestinal tissue to identify both development and NEC-associated changes. This study not only demonstrates ISC subpopulation recovery following therapeutic intervention with hPSC-EVs, but also identify two cellular mechanisms of ISC disruption in NEC, loss of OLFM4+ cells and loss of the normal HOPX+ cell expansion. Future studies will continue to investigate other rISC makers, such as mTert, DCLK1, LRIG1, and CLU, to further understand ISC expression throughout early developmental stages. Overall, our study provides a starting foundation for understanding how ISC development in the preterm infant influences NEC susceptibility and reveals potential novel therapeutic targets to improve neonatal health outcomes.

## Supplemental Materials

**Supplemental Figure 1.**
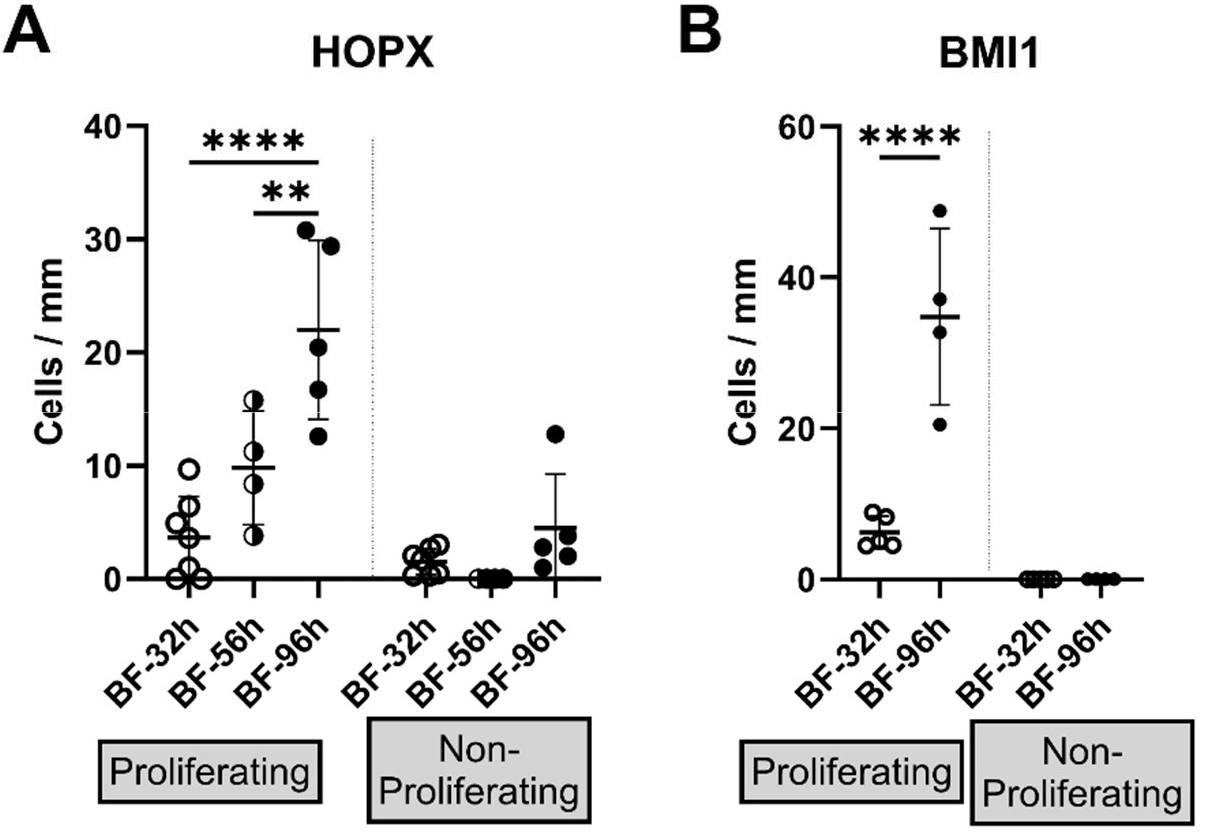
(A) Proliferating HOPX+ cells are significantly increased throughout early postnatal development timepoints in BF ileum, while non-proliferating HOPX+ cells demonstrate no significant increases. (B) Proliferating BMI1+ cells significantly increased from 32 hours to 96 hours, while non-proliferating BMI1+ cells remained undetectable across early postnatal development. Quantifications are expressed as mean ± standard deviation of cell counts normalized per length of tissue analyzed. Samples sizes for each analysis are denoted in dot scatter plots, with n=4-7 animals per group. \*\*\*\**p<*0.0001.

**Supplemental Figure 2.**
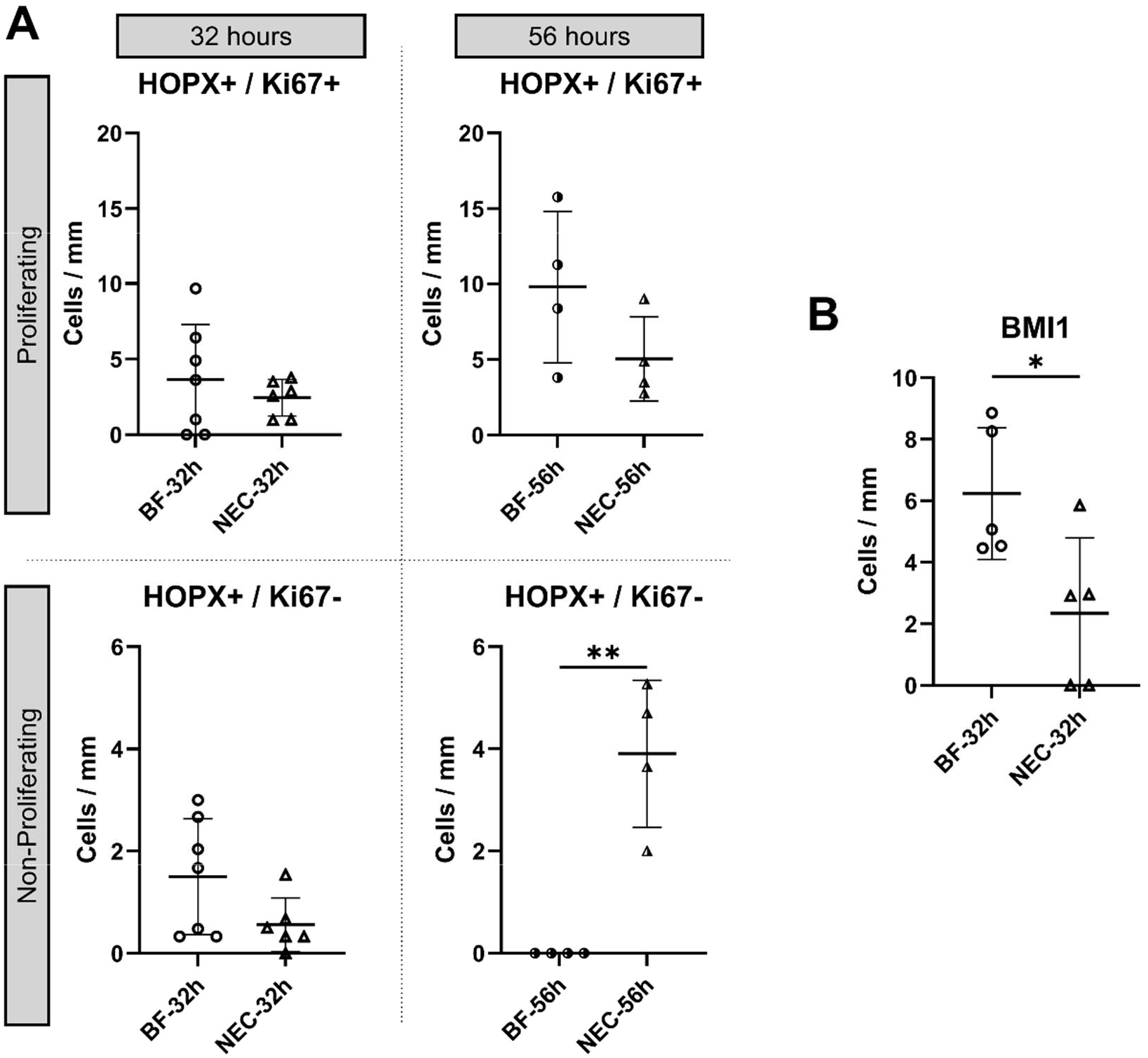
(A) Proliferating HOPX+ cells (dual labeled HOPX+/Ki67+) demonstrated no significant changes between BF and NEC at 32 hours or 56 hours. Non-proliferating HOPX+ cells (single labeled HOPX+/Ki67-) demonstrated no changes between BF and NEC at 32 hours but increased significantly in NEC at 56 hours. (B) BMI1+ cells demonstrate a significant decrease in NEC as compared to BF at 32 hours. Quantifications are expressed as mean ± standard deviation of cell counts normalized per length of tissue analyzed. Samples sizes for each analysis are denoted in dot scatter plots, with n=4-7 animals per group. \**p<*0.05, \*\**p<*0.01.

## Contributions

Study conception and design: M.A.E. and V.W.; Data acquisition: M.A.E., S.P., A.G., M.E.; Data analysis and interpretation: M.A.E and V.W.; Writing and editing the manuscript: all authors.

